# Task-irrelevant stimuli boost phasic pupil-linked arousal but not memory formation

**DOI:** 10.64898/2025.12.23.696068

**Authors:** J. Hebisch, P. Van Puyenbroeck, L. Schwabe, J.W. de Gee, T.H. Donner

## Abstract

Brainstem arousal systems including the locus coeruleus noradrenaline system, respond transiently to behaviorally relevant events. Locus coeruleus activity also drives dilations of the pupil, which are often observed during cognitive tasks. The strength of pupil responses during encoding of stimulus material predicts the success of its later retrieval, which might reflect the impact of noradrenaline on synaptic plasticity and memory formation. The pupil also dilates in response to task-irrelevant sounds, which could therefore serve as a valuable tool for investigating causal effects of phasic, pupil-linked arousal on cognition. Here, we evaluated whether task-irrelevant white noise sounds affect memory formation and memory-based decisions. These sounds were played before, during or after the presentation of memoranda (images or spoken words). Memory success was measured in recognition and free recall tasks the day after. Trial-to-trial variations in the amplitude of pupil dilations during word encoding without task-irrelevant sounds predicted memory success. Task-irrelevant white-noise sounds also robustly dilated the pupil but did not improve memory formation for the words or the images. We conclude that pupil-linked arousal processes triggered by task-irrelevant sounds differ from those recruited endogenously during memory formation, for example in states of increased emotionality or attention.

## Introduction

Not all memories are remembered with equal ease. Prioritizing the encoding and consolidation of some memories over others enables the brain to store only what will likely be relevant in the future (Richter-Levin & Akirav, 2003; Schacter & Addis, 2009; Shohamy & Adcock, 2010). For example, memories encoded under high emotionality are generally retrieved better (Bergt et al., 2018; Richter-Levin & Akirav, 2003). Mounting evidence suggests that the prioritization of memory formation by behavioral relevance may be mediated by the arousal systems of the brainstem (Clewett et al., 2017; Clewett et al., 2018; Mather & Sutherland, 2011), in particular the locus coeruleus (LC) noradrenaline (NA) system. With its widely distributed projections to the forebrain and impact on its target circuits, the LC-NA system is in an ideal position to control network states (Schwarz & Luo, 2015) as well as synaptic plasticity in the cerebral cortex (Bear & Singer, 1986; Jordan & Keller, 2023; Seol et al., 2007) and hippocampus (Bliss & Collingridge, 1993; Hagena & Manahan-Vaughan, 2025; Sonneborn & Greene, 2021), which is, in turn, required for memory formation (Dringenberg, 2020; Goto et al., 2021; Martin et al., 2000).

The LC-NA system and other components of the brain’s arousal system activate transiently during cognitive tasks (Bouret & Sara, 2005; Breton-Provencher et al., 2022; Breton-Provencher & Sur, 2019; de Gee et al., 2017), which also drives dilations of the pupil (Joshi et al., 2016; Maheu et al., 2025). Thus, pupil responses under conditions of constant luminance have often been used as a marker of transient (“phasic”) arousal responses (Joshi & Gold, 2020). Emotional stimuli evoke a larger pupil response during encoding than neutral stimuli (Bergt et al., 2018; Goldinger & Papesh, 2012; Lempert et al., 2015). The amplitude of these pupil responses, in turn, predict subsequent memory success (Bergt et al., 2018; Kucewicz et al., 2018). While such results are consistent with a role of brainstem arousal systems in memory formation, these findings are correlative in nature.

Task-irrelevant sounds have emerged as a promising non-invasive method for causally driving pupil (Cronin et al., 2023; Hebisch et al., 2024; Joshi et al., 2016; Nassar et al., 2012; Petersen et al., 2017; Tona et al., 2016) as well as LC responses (Joshi & Gold, 2022; Joshi et al., 2016). Because biases in perceptual decisions are affected by optogenetic LC stimulation in mice (Breton-Provencher et al., 2022) and pharmacological catecholamine boost in humans (de Gee et al., 2025), we previously tested the effects of task-irrelevant sounds on perceptual decision-making (Hebisch et al., 2024). While the task-irrelevant sounds evoked reliable pupil dilations, they did not affect biases or any other aspect of perceptual decision-making (Hebisch et al., 2024). These findings cast doubt on the suitability of task-irrelevant sounds as a tool for effectively modulating decision-making, but do not necessarily imply a failure to trigger activity in the brainstem arousal system. For example, it is possible that white noise stimuli drive responses in LC sub-populations whose projections do not affect perceptual decision-making (Breton-Provencher et al., 2022).

In the present study, we tested the possibility that task-irrelevant sounds boost memory encoding success, rather than affecting decision-making. Given the apparently modular organization of the LC (Schwarz & Luo, 2015) and the established role of noradrenergic arousal in synaptic plasticity, such a specific effect is plausible and would offer valuable insights into the function and specificity of brainstem arousal systems. To this end, we asked participants to encode either spoken words or images in memory, followed by free recall and a recognition memory test on the next day. Task-irrelevant white noise sounds were played at different time points during encoding as well as the recognition task. If task-irrelevant sounds effectively drive LC responses, they should dilate the pupil and, critically, boost memory success for the images or words that are paired with the sounds during encoding. The (“old” versus “new”) recognition task enabled us to test for an effect of task-irrelevant white noise sounds on memory decision bias, which, just as perceptual decision bias (de Gee et al., 2017; Hebisch et al., 2024; Nuiten et al., 2025), negatively correlates with the amplitude of task-evoked pupil responses (de Gee et al., 2020).

## Results

Participants encoded 150 greyscale neutral images and 60 neutral words on day 1 of the experiment (**Fig. 1A**; see Methods). Task-irrelevant sounds were played on 40% (image task) or 27% (word task) of randomly selected trials with an onset time before, during (only image task) or after stimulus presentation (randomly selected). Task-irrelevant sounds consisted of 3 s of white noise at 75 dB, which evoked robust pupil dilations in our previous work (Hebisch et al., 2024). Right after encoding, participants were asked to freely verbally recall as many stimuli as they could. Day 2 of the experiment (24 hours later) started with another free verbal recall of the stimuli, followed by a recognition task (**Fig. 1D**). In the latter, we presented 300 images and 120 words, half of which were repeats from day 1 and half of which were new. Participants were asked to report whether they thought the stimulus was “old” or “new” by pushing one of two buttons. They then had to indicate their confidence in the decision on a scale from 1 to 4. During the recognition task, task-irrelevant sounds were again played on 40% of randomly selected trials, either before or during stimulus presentation.

**Figure 1.**
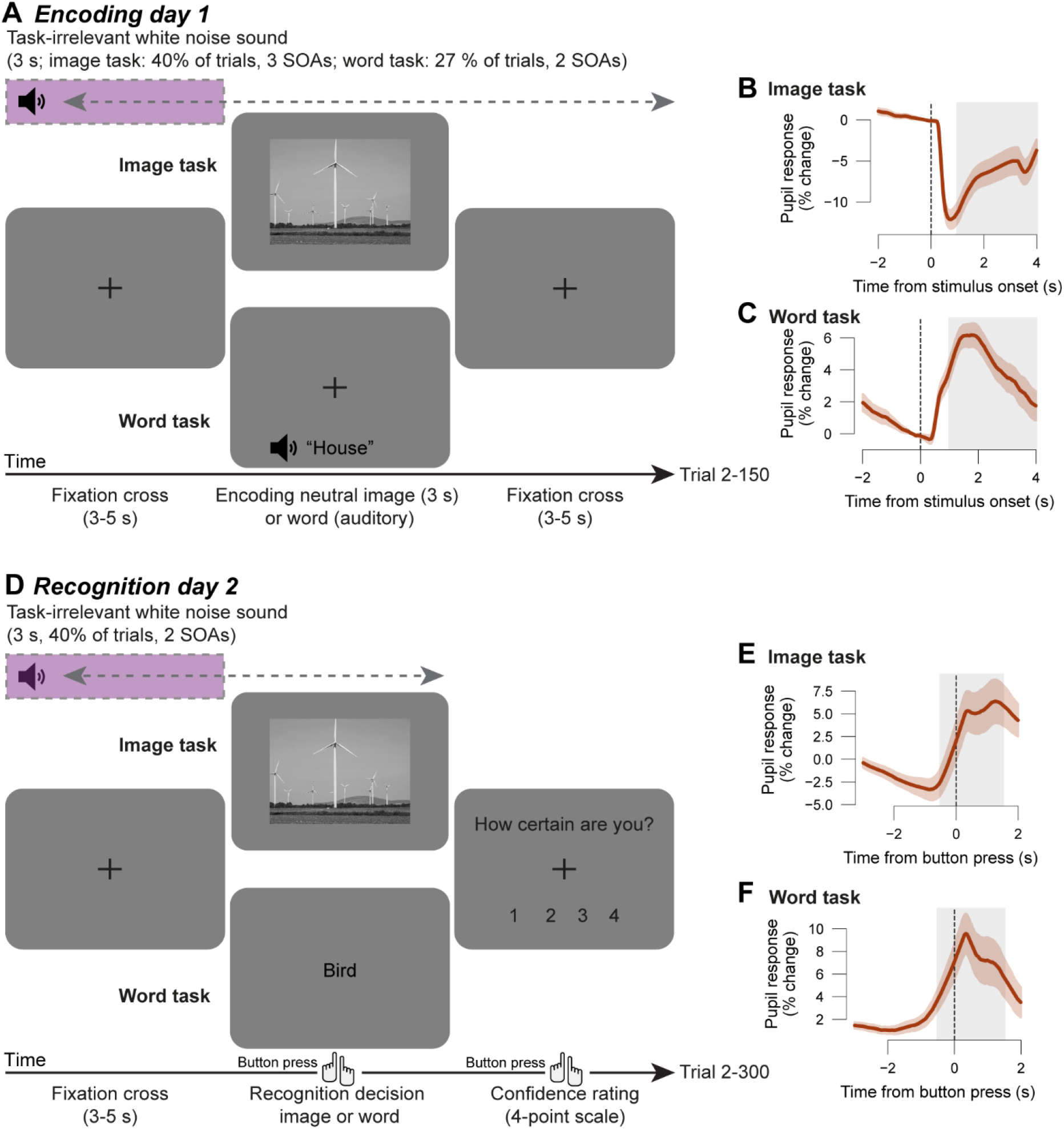
Behavioral tasks and pupillometry. **(A)** Schematic sequency of trial events during encoding (day 1). The image task comprised 150 greyscale images (mean luminance matched with grey background), and the word task comprised 60 words (presented via headphones). The task-irrelevant, auditory white noise sound appeared before, during, or after image presentation on a total 40% of trials, and before or after word presentation on a total of 27% of trials. **(B)** Average pupil response time course during encoding, locked to image onset. Lines, group average; orange shading, SEM across participants; grey shading, time window used for quantifying task-evoked pupil responses. **(C)** Same as (B) but for words task. **(D)** The recognition task (day 2) consisted of 300 images or 120 visually presented words respectively and task-irrelevant sounds were played before or during stimulus presentation on a total of 40% of trials. **(E-F)** As (B-C) but for pupil responses during recognition task, locked to choice (button-press).

The mean task-evoked pupil responses on trials without any task-irrelevant sounds replicated effects previously reported for the encoding task (Bergt et al., 2018): pupil constriction locked to image presentation (**Fig. 1B**) and dilation locked to word presentation (**Fig. 1C**). In the recognition tasks for both stimulus domains, we found pupil dilations around the time of the button press (**Fig. 1E-F**), likely reflecting phasic task engagement (de Gee et al., 2017; de Gee et al., 2014; de Gee et al., 2020).

### Evoked pupil responses during word encoding predict memory success

A previous study found a correlation between the amplitude of pupil responses during stimulus encoding and subsequent memory performance (Bergt et al., 2018). Such an effect was observed for both emotional words and images, but when stimuli were neutral, it was only observed for words, not for images (Bergt et al., 2018). We replicated the effects for neutral stimuli in our current dataset, which did not contain emotional stimuli.

We quantified pupil response amplitude as the mean response over a window of 1-4 s following stimulus onset on trials without task-irrelevant sounds (see Methods). No effects related to memory success were evident for the image task (BF_10_ ranging from 0.423 to 0.913; **Fig. 2A**,**C**; pupil response time courses split by memory success, **Fig. S1**). However, words that were remembered in free recall (day 2) elicited a larger pupil response during encoding than words that were later forgotten (t=1.759, p=0.047, one-sided t-test; BF_10_=1.706; **Fig. 2D**). Splitting pupil responses based on whether the stimuli were successfully remembered in the recognition task yielded a similar effect, which was only trending (t=1.555, p=0.068, one-sided t-test; BF_10_=1.303; **Fig. 2B**; see **Fig. S2** for immediate free recall). In sum, we found a weak, but significant relationship between pupil responses during encoding of neutral words and later memory success that was qualitatively similar to previously reported results (Bergt et al., 2018).

**Figure 2.**
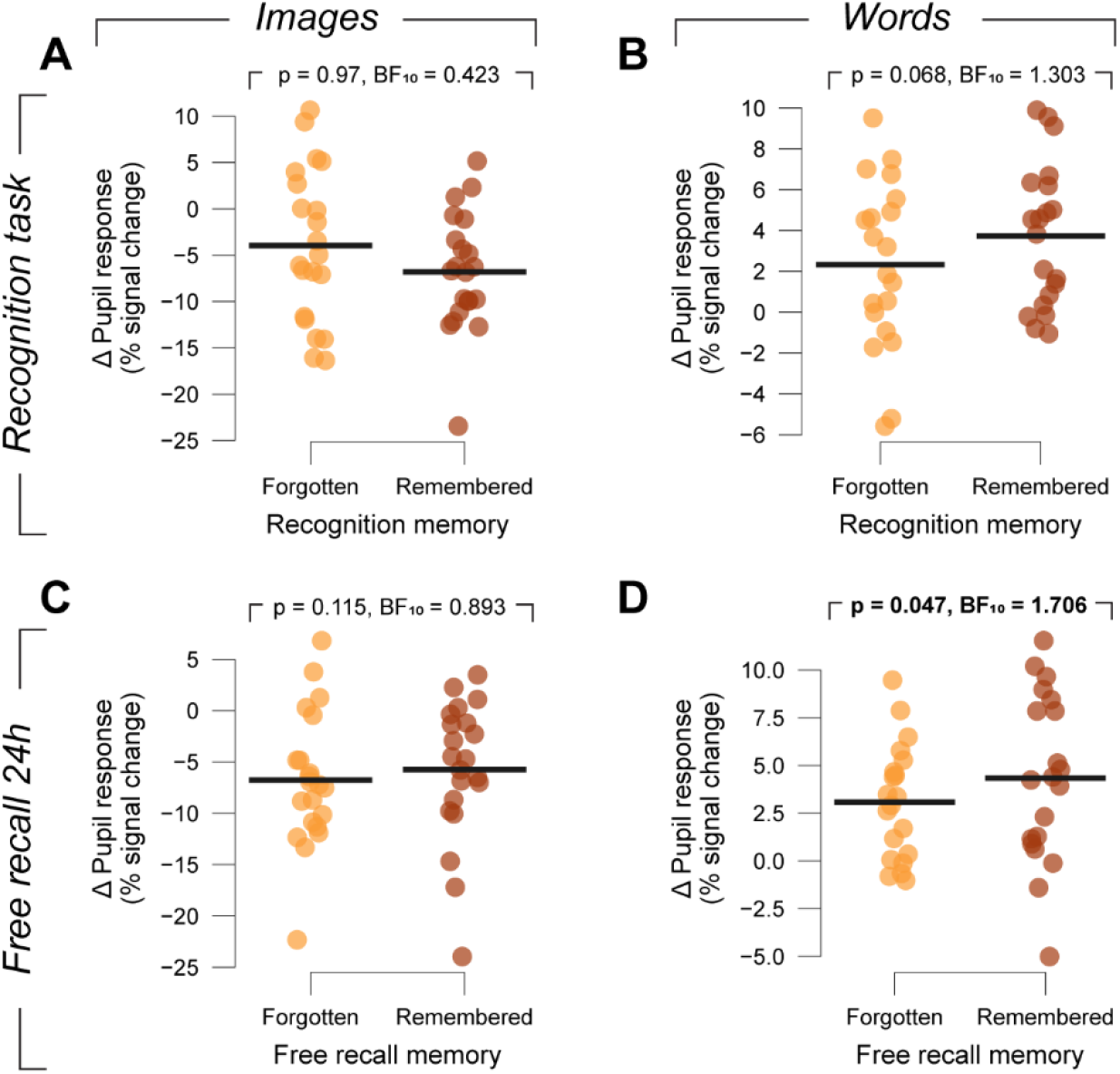
Amplitude of responses during encoding sorted by subsequent memory performance. **(A)** Individual trial-average pupil response amplitudes during encoding (collapsed across an interval of 1-3 s from image onset) on trials with no task-irrelevant sounds. Trials are sorted by whether images were recognized (remembered) or missed (forgotten) in the recognition task on day 2. Black bars, group average. **(B)** As (A), but for words. **(C, D)** As (A) and (B), but for images and words that were recalled vs. not recalled 24h later.

### Task-irrelevant sounds drive pupil dilations

We next isolated the pupil response evoked by the task-irrelevant sound by computing differential response time courses between trials with and without task-irrelevant sounds (see Methods). For each stimulus onset asynchrony (SOA) between task-irrelevant sound and memorandum onset, the differential time course showed clear dilations for roughly 4 seconds (i.e. until 1 s after the sound terminated, **Fig. 3A-D**), peaking at about 10% signal change.

**Figure 3.**
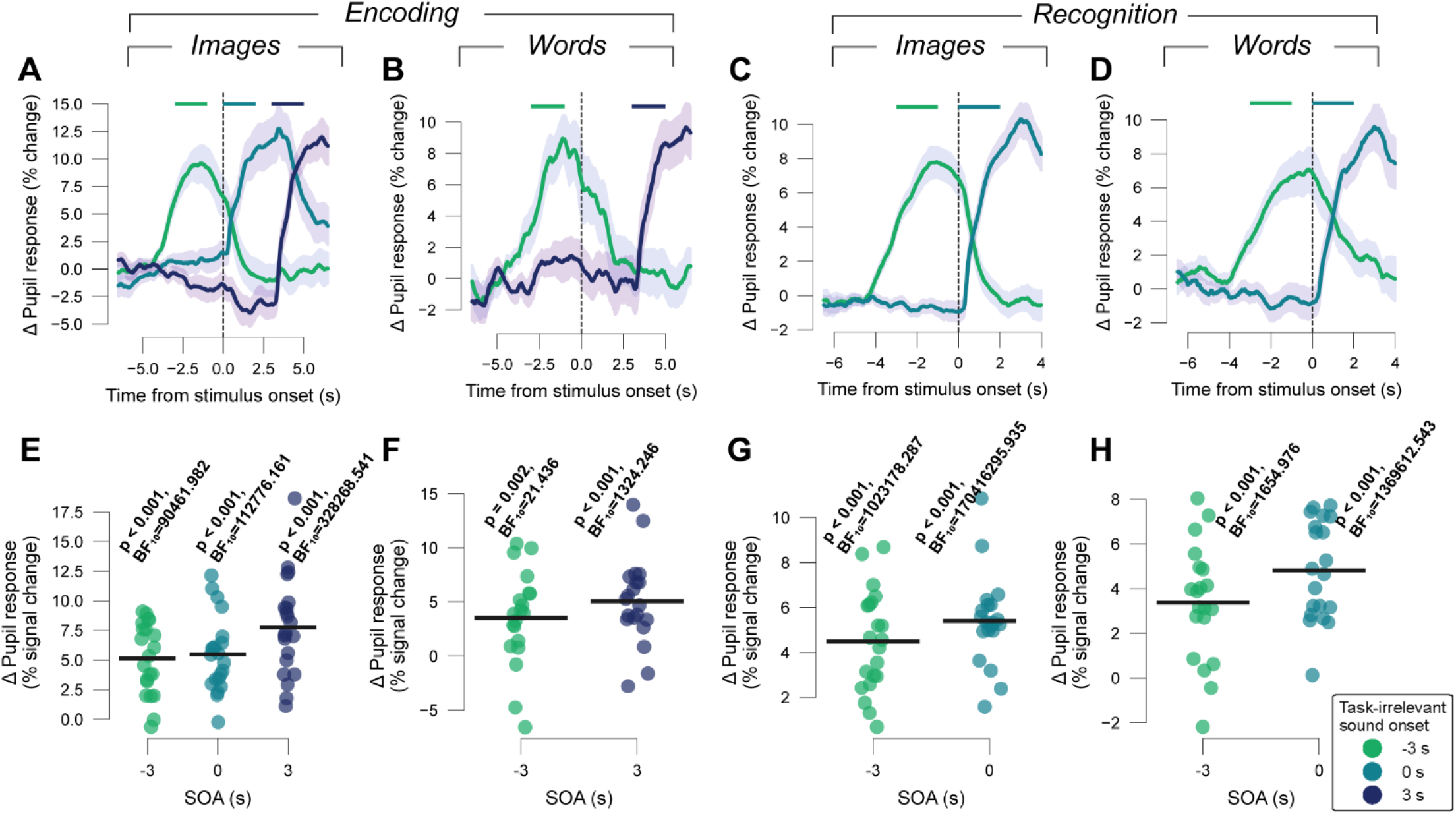
Robust pupil responses to task-irrelevant sounds. **(A)** Differential pupil response time courses between trials with or without task-irrelevant sounds in the image encoding task. Time courses were baseline-corrected with mean value from interval from 3.5 to 3 s prior to stimulus onset. Shading, SEM across participants. Top bars, intervals used for quantifying task-irrelevant sound-evoked pupil responses for the respective sound onsets. **(B)** As (A) but for word encoding task. **(C-D)** As (A-B), but for the recognition tasks. **(E)** Individual amplitudes of sound-evoked pupil responses in image encoding task. Black bars, group average. P and BF_10_, p-values and Bayes factors (two-sided t-tests against zero). **(F)** As (E) but for the word encoding task. **(G-H)** As (E-F), but for the Recognition task.

We quantified the amplitude of the sound-evoked pupil response by averaging the differential response time courses across 2 seconds following sound onset. Most participants exhibited reliable positive (i.e., dilatory) effects (**Fig. 3E-H**), with highly significant group-level effects of task-irrelevant sounds for each task (main effects of sound condition: image encoding task: F_(3,60)_=33.296, p<0.001; word encoding task: F_(2,38)_=15.305, p<0.001; image recognition task: F_(2,40)_=72.747, p<0.001; word recognition task: F_(2,38)_=34.415, p<0.001; repeated measures ANOVAs). This was further supported by Bayesian t-tests for the different SOAs (see Methods; range of BF_10_: from 21.436 to 170416295.935). These results replicate our previous research that comprehensively quantified effects of task-irrelevant white noise sounds of different durations (including the 3 s used here) on pupil responses in the context of a perceptual decision task (Hebisch et al., 2024).

### Task-irrelevant sounds do not boost memory formation

Despite robustly driving pupil dilations, the task-irrelevant sounds had no detectable effect of boosting memory formation (**Fig. 4**). We observed no evidence for an effect of sound condition (no sound, sound before, during or after stimulus presentation in the encoding task) on the success of recognition (image task, **Fig. 4A**: F_(3,60)_=0.172, p=0.915; word task, **Fig. 4B**: F_(2,38)_=1.674, p=0.201) or free recall (image task, **Fig. 4C**: F_(3,60)_=1.719, p=0.173; word task, **Fig. 4D**: F_(2,38)_=2.035, p=0.145; see **Fig. S2** for immediate free recall on day 1). The differences in recognition accuracy between trials with and without task-irrelevant sounds were not likely different from zero for any SOA (BF_10_ range from 0.228 to 0.476). The same was true for memory accuracy on the free recall (BF_10_ range from 0.229 to 0.624) that took place right before the recognition task, with one notable exception: words which were followed by the task-irrelevant sound were recalled significantly less often than those without task-irrelevant sounds during encoding (t=-2.971, p=0.008, BF_10_=6.268). This may result from an interfering effect of the sounds after word presentation on memory encoding. The sound condition also did not impact confidence ratings in the recognition task (image task: F_(3,60)_=0.276, p=0.842, SOA -3 s: BF_10_=0.296, SOA 0 s: BF_10_=0.228, SOA 3 s: BF_10_=0.233; word task: F_(2,38)_=0.217, p=0.806, SOA -3 s: BF_10_=0.278, SOA 3 s: BF_10_=0.233). In sum, task-irrelevant sounds during encoding do not boost memory formation.

**Figure 4.**
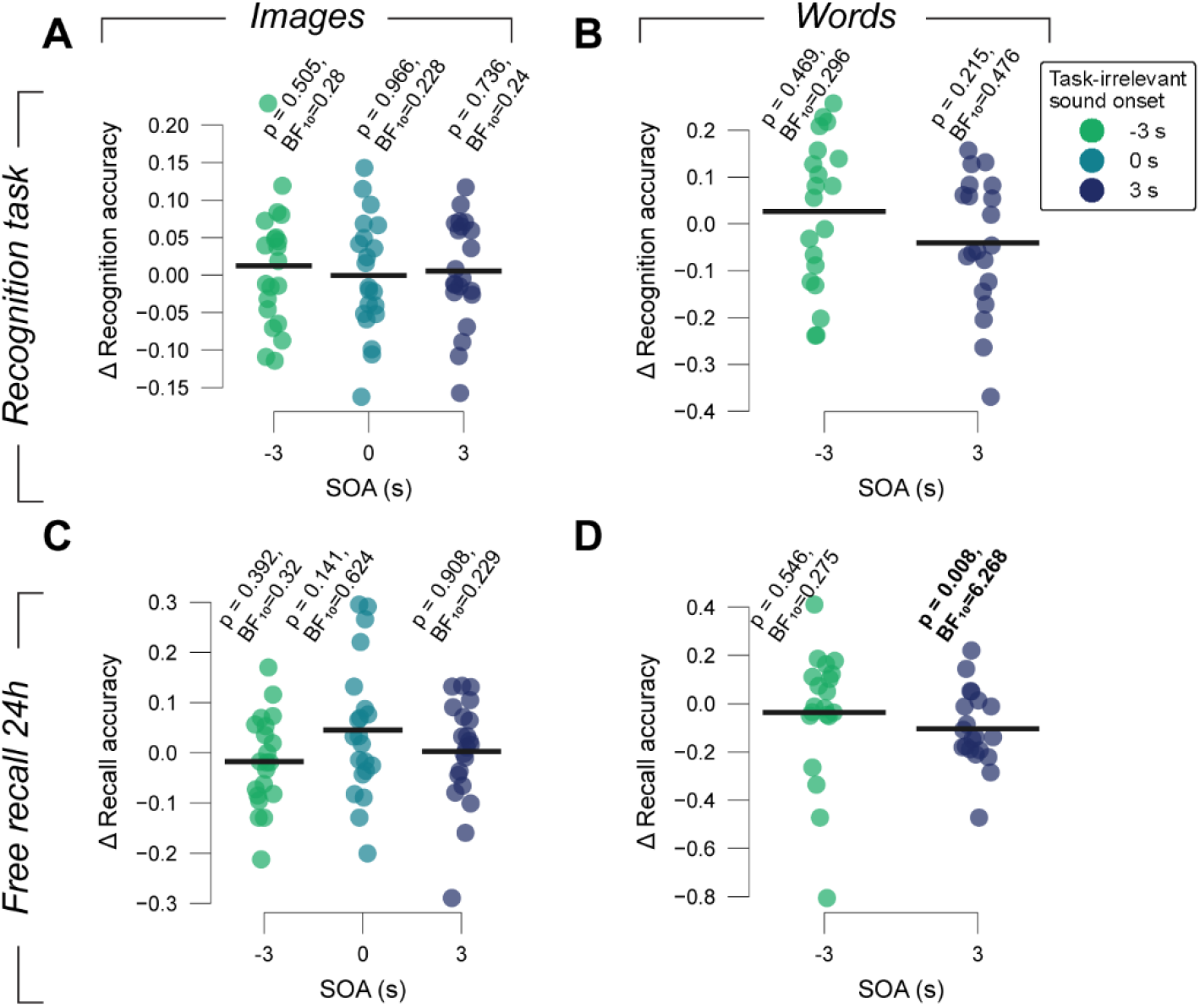
Difference in memory performance due to task-irrelevant sounds in encoding. **(A)** Difference of recognition accuracy between trials with and without task-irrelevant sounds (played in the encoding task) for each individual, sorted by sound SOA. Black bars, group average. P and BF_10_, p-values and Bayes factors for t-tests against zero. **(B)** As (A), but for word tasks. **(C-D)** As (A-B), but for accuracy during free recall on day 2.

### Task-irrelevant sounds also do not affect recognition decision

Recognition task performance does not only depend on the quality of memory encoding, but additionally on the (memory-based) decision process involved in deciding whether an image is old (i.e. familiar) or new (unfamiliar; Ratcliff, 1978). Previous work had established a negative relationship between the amplitude of pupil dilations during recognition decisions and the (commonly conservative, favoring “new” responses) bias in those decisions (de Gee et al., 2020). We replicated the above effect for the trials without task-irrelevant sound during the recognition task (**Fig. S3**). The task-evoked pupil responses around choice report varied substantially from trial to trial, with dilations on the most trials, but even robust constrictions from baseline on some others (**Fig. S3A-B**). The amplitude of the evoked pupil response correlated negatively with signal detection-theoretic measure of bias – decision criterion c – in the (image) recognition decision: the larger the pupil response, the more negative c, so the larger the tendency to report that an image was “old” (**Fig. S3C**, group average correlation coefficient r_s_=-0.24, t=-3.1, p=0.006). This effect was not observed on the word recognition task (**Fig. S3D**, group average correlation coefficient r_s_=-0.172, t=-1.738, p=0.098). While the relationship of task-evoked pupil responses with bias was, for image recognition, approximately linear, the pupil responses exhibited a non-linear relation to recognition sensitivity (signal detection-theoretic index d’; **Fig. S3E-F**), captured by a second-order polynomial (polynomial regression, see Methods). This inverted U-shape means that intermediate task-evoked responses went along with maximal sensitivity for both recognition tasks. Together, these results illustrate two distinct (monotonic versus non-monotonic) effects of phasic, arousal responses on decision bias and sensitivity, in line with previous work (de Gee et al., 2020).

Despite these correlations between task-evoked pupil responses and decision performance (**Fig. S3**) as well as the robust pupil responses to task-irrelevant sounds during the recognition task (**Fig. 3C-D, G-H**), task-irrelevant sound conditions per se had no effect on recognition behavior (**Fig. 5**). We found no evidence for a modulation of decision bias (c) through task-irrelevant sounds in either the image or the word recognition tasks (image task: F_(2,40)_=0.616, p=0.545, SOA -3 s: BF_10_=0.295, SOA 0 s: BF_10_=0.244; word task: F_(2,38)_=0.87, p=0.427, SOA -3 s: BF_10_=0.458, SOA 0 s: BF_10_=0.236; **Fig. 5A-B**). The same was true for sensitivity (d’) in the image task (image recognition task: F_(2,40)_=0.893, p=0.417; BF_10_ range from 0.228 to 0.387, **Fig. 5C-D**). Solely in the word recognition task, there was an effect of task-irrelevant sound (word recognition task: F_(2,40)_=4.082, p=0.025, repeated measures ANOVA), where task-irrelevant sounds played before word presentation reduced sensitivity (t=-3.139, p=0.005, BF_10_=8.586, **Fig. 5D**), again in line with distracting effect of the sounds. We also found no effect of task-irrelevant sounds on reaction times (image recognition task: F_(2,40)_=0.874, p=0.425, SOA -3 s: BF_10_=0.292, SOA 0 s: BF_10_=0.517; word recognition task: F_(2,38)_=1.588, p=0.218, SOA -3 s: BF_10_=0.513, SOA 0 s: BF_10_=0.236; **Fig. 5E-F**) or confidence ratings (image recognition task: F_(2,40)_=0.775, p=0.468, SOA -3 s: BF_10_=0.539, SOA 0 s: BF_10_=0.316; word recognition task: F_(2,38)_=1.47, p=0.243, SOA -3 s: BF_10_=1.274, SOA 0 s: BF_10_=0.27).

**Figure 5.**
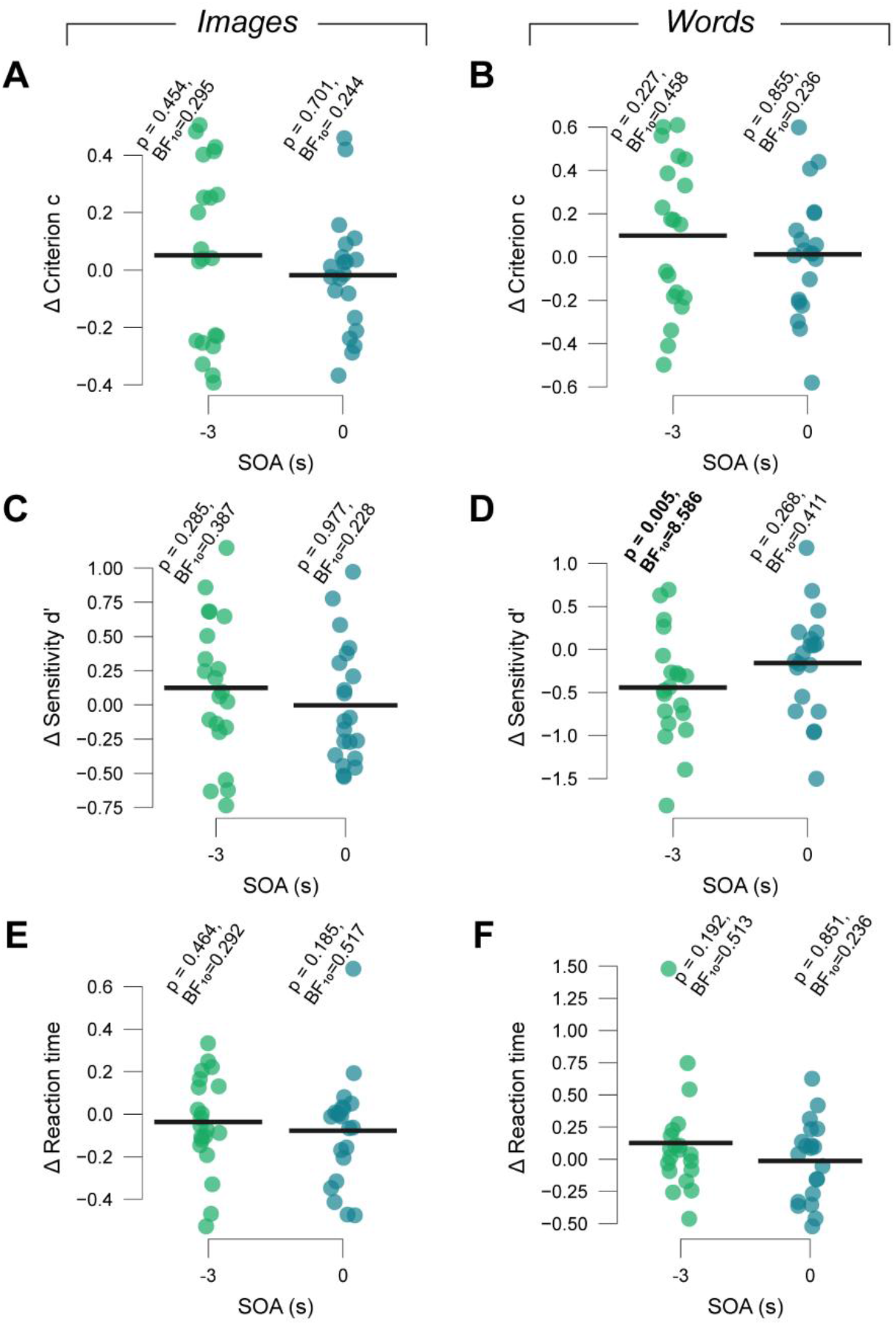
Behavioral effects of task-irrelevant sounds during recognition tasks. **(A)** Participant-wise difference of recognition decision bias (criterion c) on task-irrelevant sound trials versus trials with no task-irrelevant sound for the respective sound onset asynchrony with respect to stimulus onset. Black bars, group average. P and BF_10_, p-values and Bayes factors for t-tests against zero. **(B)** As (A), but for word recognition task. **(C-D)** As (A-B) but for recognition performance (sensitivity d’). **(E-F)** As (A-B) and (C-D), but for reaction time.

## Discussion

Previous correlative studies have pointed to a key role for phasic, pupil-linked arousal in the prioritization of some memories over others (Bergt et al., 2018; Clewett & Murty, 2019; Clewett et al., 2018; Kucewicz et al., 2018; Mather & Sutherland, 2011), in line with a role of NA, and other neuromodulators controlling central arousal states, in cortical plasticity. We tested if such prioritization of memories could be causally controlled by driving phasic pupil-linked arousal via task-irrelevant sounds. We found that words that were later remembered tended to be associated with larger pupil dilations during encoding compared to later forgotten words. Task-irrelevant sounds during encoding, however, did not improve later memory success, despite evoking robust pupil dilations. In other words, the predictive value of pupil dilation per se for memory success found in previous work (Bergt et al., 2018) and replicated here, does not hold for all contexts.

There is strong evidence that phasic LC activation boosts synaptic plasticity in the hippocampus (Hagena & Manahan-Vaughan, 2025) and cerebral cortex (Bear & Singer, 1986; Jordan & Keller, 2023; Seol et al., 2007) which is critical for memory formation (Dringenberg, 2020; Martin et al., 2000). For example, NA release shapes long-term potentiation and depression in the hippocampus (Bliss & Collingridge, 1993; Harley, 2007; Sonneborn & Greene, 2021). Had our task-irrelevant sounds led to LC activity of enough extent, we would have expected effects on memory success. Loud sounds, such as the ones we used here, drive LC activity and dilate the pupil (Grant et al., 1988; Joshi & Gold, 2022; Joshi et al., 2016). However, such sounds evoke even larger responses in the inferior colliculi (Joshi et al., 2016), which are part of the early auditory pathway (Ades & Brookhart, 1950), sufficient to dilate the pupil (Joshi et al., 2016), but play no obvious role in memory formation. Stronger drive of inferior colliculi than LC could explain the observed combination of robust pupil dilations without concomitant memory improvement under task-irrelevant sounds.

Bypassing the LC by task-irrelevant sounds (or triggering negligible LC responses) is only one possible explanation of our current set of results. It is also possible that the rapid, sound-evoked arousal boost does not match arousal dynamics that naturally regulate memory formation, for example due to the typically longer time scales of emotional dynamics (Tambini et al., 2017). Yet another idea is that the sounds did drive LC neurons, but a different subpopulation of those neurons than the ones required for memory formation, for example due to different patterns of projections to the forebrain (Breton-Provencher et al., 2022; Poe et al., 2020; Totah et al., 2018). In light of analogous results of the sound manipulation in the context of perceptual decision-making (Hebisch et al., 2024), the lack of effect of white noise sounds on behavior seems to generalize across domains of cognition. We, therefore, consider it likely that such sounds are generally not sufficient to drive the LC neuronal populations necessary for measurable effects on cognitive behavior.

Our findings are overall in line with another study that found an effect of different types of task-irrelevant sounds on pupil dilation without any effect on face recognition performance (Cronin et al., 2023). However, as the authors of this study concluded, the lack of effect on recognition performance could have been due to misaligned timing: in their experiment, sounds were played exclusively before stimulus onset. The systematic manipulation of sound onset times relative to the memoranda allows us to rule out this explanation here. In addition, we extended that previous work by testing memory 24 h after encoding instead of in the same session, which (i) established an effect of task-evoked pupil dilation on long-term memory success and (ii) ruled out that overnight consolidation may culminate in a beneficial impact of task-irrelevant sounds on memory.

Notably, we found a negative effect of the sounds being played after auditory presentation of stimulus in the word encoding task on free recall accuracy on day 2. This points to an interference of auditory encoding through the task-irrelevant sounds. Interference shortly after encoding may play a central role in forgetting (Lewandowsky et al., 2009). The fact that we did not observe this for the image encoding may be caused by the difference in modality between the task-irrelevant stimulus and the stimulus to be encoded.

We also studied the effects of the task-irrelevant sounds during the recognition task as well as of trial-to-trial variations in task-evoked pupil responses. Regarding the former, we found no clear impact of task-irrelevant sounds on decision behavior, aligning with findings in perceptual decisions (Hebisch et al., 2024). Regarding the latter, we found a negative, linear relationship between recognition task-evoked pupil response magnitude and decision bias (i.e. old/new tendency) in the image task, but not the word task. This further underscores the robustness of earlier findings of this relationship between phasic pupil-linked arousal in decision-making both in perceptual decision-making and decisions from memory (de Gee et al., 2017; de Gee et al., 2020; Hebisch et al., 2024; Nuiten et al., 2025). This effect may reflect an influence of arousal on the weighting of priors relative to incoming evidence in the decision process (Bouret & Sara, 2005; Dayan & Yu, 2006; de Gee et al., 2017; Krishnamurthy et al., 2017), whereby the evidence is here sampled from memory (Shadlen & Shohamy, 2016). Further, task-evoked pupil responses in both the image and the word task, exhibited an inverted U-shaped relationship with decision sensitivity. This adds to the vast literature around the so-called Yerkes-Dodson law (Teigen, 1994; Yerkes & Dodson, 1908), describing maximal cognitive performance during mid-level arousal states (Beerendonk et al., 2024; McGinley et al., 2015). However, we point out that this literature primarily describes behavioral correlates of slowly varying states of baseline arousal (Beerendonk et al., 2025; McGinley et al., 2015; Nuiten et al., 2025), rather than correlates of the rapid, task-evoked responses for which we here found this inverted-U relationship.

The ability to non-invasively manipulate phasic LC activity (and the activity of other brainstem systems) could profoundly advance the mechanistic understanding of the relationship between neuromodulation and cognitive processes relying on synaptic plasticity, such as memory and learning. The lack of impact of task-irrelevant sounds on memory-related behavior, that we found here, suggests that they are not suitable for this aim.

## Methods

### Participants

23 healthy volunteers (11 females, 12 males; age range, 20-33 y) participated in this experiment. All participants had normal or corrected-to-normal eyesight without wearing glasses. Participants were instructed not to drink any beverages containing any caffeine up to two hours prior to the sessions.

All participants gave written informed consent and were remunerated by the hour (15 €/h). The experiment was conducted in accordance with the Declaration of Helsinki. It was approved by the ethics committee of the Faculty of Psychology and Human Movement Sciences at the University of Hamburg. Two participants were excluded due to a low number of valid trials (<250) after removing trials with more than 20% missing pupil data (see analysis of pupil responses) or reaction times below 200 ms or longer than 8 s. This resulted in a sample size of 21 participants for the image tasks and 20 participants in the words task.

### Procedure

The experiment consisted of two consecutive sessions that took place on subsequent days, with a delay of about 24 h. Each session lasted approximately 2 h. In both sessions, participants were asked to sit at a desk resting their head on a chin rest at 50 cm distance of the VIEWPixx monitor (1920 by 1080 pixels) with a refresh rate of 100 Hz. Luminance levels were kept stable throughout the whole task. The experiment took place in a dark room (0 Lux). The screen color was kept stable throughout the experiment tasks at an average RGB value of [128, 128, 128]. The first session consisted of an image encoding task, followed by an immediate free recall, and then a word encoding task, followed by a free recall as well. The second session consisted of: the free recall of images, the free recall of words, an image recognition task, and finally a word recognition task. This experimental design was adapted from the one used by Bergt et al. (2018). Stimulus sets contained neutral images and words (written and audio). These included stimuli used by Bergt et al. (2018) and images from the THINGS database (Hebart et al., 2019). Images were presented in greyscale and were modified using the Pillow toolbox in python (Clark, 2015) to have the equal average luminance, which was the same as the grey background.

### Apparatus

The experiment was controlled by a personal computer using the MATLAB Psychophysics Toolbox (Brainard, 1997). Auditory stimuli were played using AKG K72 headphones.

The gaze direction and pupil size of participants’ right eyes were tracked using an Eyelink 1000 (SR Research, Osgoode, Ontario, Canada). The eye data was sampled at a rate of 1000 Hz and an average spatial resolution of 15 to 30 min arc. At the beginning of each task, the eye tracker was calibrated. Obtaining data from one eye is standard practice as there is no evidence pointing to differences in pupil responses between both eyes in participants without neurological disorders (Wang & Munoz, 2014).

### Encoding tasks

Trials in the image and word encoding tasks consisted of a pre-stimulus interval (3-5 s), a stimulus presentation interval (3 s) and a post-stimulus interval (3-5 s). During the pre- and post-stimulus intervals, only a fixation cross was shown on the grey background. In the image encoding task, we presented an image overlaid with a fixation cross during the full duration of the stimulus presentation interval. In total, participants saw 150 images. In the word encoding task, participants heard a word in the beginning of the stimulus presentation interval (average duration ∼1 s) while only a fixation cross was presented on the screen. In total, participants heard 60 words.

On a fraction of trials, a task-irrelevant sound (auditory white noise, 75 dB, 3 s duration) occurred on one of multiple possible stimulus onset asynchronies (SOAs). These trials were randomly selected under the constraint that the same number of sounds are played at each SOA. In the image encoding task, this task-irrelevant sound was played on 40% of trials with an onset either in the pre-stimulus interval (3 s prior to image onset), at the same time as image onset or in the post-stimulus interval (3 s after image onset). In the word encoding task, it was played on 27% of trials either in the pre-or the post-stimulus interval.

### Free recall tasks

In the free recall task, participants were asked to verbally recall as many stimuli as possible. For the image recall task, this meant that participants first mentioned as many details about each image as they could describe with ease. If the experimenter was unsure which image was described back to them, they asked to describe the image in more detail until they could either identify the image, or the participant could not go into more detail. During this task, participants sat behind a curtain to prevent effects of non-verbal communication between participants and experimenter.

### Recognition tasks

In the recognition tasks, each trial consisted of a pre-stimulus interval (3-5 s), a decision interval (terminated by button press), and a confidence rating interval (terminated by button press). The pre-stimulus intervals consisted only of a fixation cross shown on the grey background. During the decision interval, a stimulus was presented visually and participants were requested to indicate whether this stimulus had been presented on the day before or not by button press on a computer keyboard (button “1” for old stimuli and “0” for new ones). In the confidence rating interval, participants were asked to indicate how certain they were of their answer on a scale from 1 (“not certain”) to 4 (“very certain”). In a randomized order, 300 images were presented in the image recognition task, and 60 words were presented in the word recognition task.

The task-irrelevant sounds (same duration and intensity as during encoding) were played on 40% of trials in the recognition tasks. They occurred either in the pre-stimulus interval (SOA -3 s with regard to stimulus onset) or started at the same time as the stimulus presentation in the decision interval. Trials were assigned randomly under the constraint that each SOA occurred the same number of times and that “old” stimuli paired with a sound in the recognition tasks were equally likely to have been paired with a sound in the encoding task compared to stimuli not paired with a sound in the recognition task.

### Analysis of pupil responses

Pupil data were analyzed using Python with a similar pipeline as in previous studies (de Gee et al., 2017; Hebisch et al., 2024).

#### Preprocessing

We used the Eyelink software for blink and saccade detection and a custom algorithm for detecting additional blinks missed by the software (van den Brink et al., 2016). Missing data (mainly due to blinks) was linearly interpolated between 200 ms before and 200 ms after the missing data. We corrected for pupil responses to blinks and saccades with the help of a double gamma function convolution (Knapen et al., 2016). Finally, the cleaned time series was converted from arbitrary size units provided by the eye tracker (based on pixel number) to percent modulation around the median value.

#### Quantification of task-evoked pupil responses

For the encoding tasks, we quantified evoked pupil responses locked to the stimulus onset (i.e. image or word onset, respectively) on trials with no task-irrelevant sound. Epochs with more than 20% of missing data were excluded from analysis. The remaining epochs were then baseline-corrected by subtracting the mean pupil size over a window of 0.5 s prior to stimulus onset. The scalar amplitudes of these task-evoked pupil responses were computed by averaging pupil response values across the interval from 1 s to 4 s after stimulus onset. This window was chosen in accordance with previous studies (Bergt et al., 2018) to account for the delay and sluggishness of the pupil response (de Gee et al., 2014; Hoeks & Levelt, 1993). The resulting amplitudes of task-evoked pupil responses during encoding were sorted according to whether the stimuli of the respective trials were subsequently remembered or forgotten, in either the free recall or the recognition tasks on day 2 (**Fig. 2**).

For the recognition tasks, we used epochs locked to the choice report (button press) on trials without task-irrelevant sounds. Again, epochs with more than 20% of missing data were excluded from the analysis, epochs were baseline-corrected by subtracting the mean pupil size over a window 0.5 s before stimulus onset, and the scalar amplitudes of these task-evoked pupil responses were calculated by averaging pupil response values in the interval -0.5 s to 1.5 s from button-press, again chosen based on previous work quantifying decision-related pupil responses (de Gee et al., 2014; Hebisch et al., 2024). We used linear regression to remove variations in pupil response due to pre-trial baseline pupil size or reaction time. The resulting amplitudes of task-evoked pupil responses during recognition were sorted into eight bins, within each participant (**Fig. S3A-B**).

#### Quantification of task-irrelevant sound-evoked pupil responses

To quantify the pupil responses evoked by the task-irrelevant sounds, we baseline-corrected epochs locked to the task-relevant stimulus onset (subtracting the mean over 0.5 s before stimulus onset), for both the encoding and the recognition tasks. Epochs with more than 20% of missing data were excluded from analysis. We then calculated the mean pupil response time courses on trials with no task-irrelevant sound and subtracted this from each pupil response time course on single trials containing a task-irrelevant sound. The resulting differential pupil epochs of trials with task-irrelevant sound were then realigned to sound onset and again baselined with the mean over a 0.5 s interval before sound onset, yielding the final sound-evoked response time courses, which were binned by SOA and then averaged. The scalar sound-evoked pupil response amplitudes were finally quantified as the mean differential pupil size over a window of 2 s following sound onset.

### Analysis of choice behavior

For all behavioral analyses, we excluded the first 10 trials of the respective task of interest due to potential effects of strong decrease in baseline pupil size in the beginning of each experimental block (compare Hebisch et al., 2024).

For analysis of effects of the task-irrelevant sound during encoding on memory performance, we computed the mean memory accuracy on immediate free recall, 24h free recall and recognition task as the fraction of correctly recalled or recognized stimuli of all stimuli presented on day 1 for the image and the word task separately. This was done separately for trials with and without task-irrelevant sounds, the former sorting by SOA of the task-irrelevant sound.

For relating trial-to-trial variations of pupil responses or the task-irrelevant sounds during recognition to behavioral performance, we computed the signal detection-theoretic indices sensitivity d’ and criterion c (Green & Swets, 1966; Macmillan & Creelman, 2005) as well as reaction time for each respective task-evoked pupil response bin or task-irrelevant sound SOA. The sensitivity reflects one’s ability to discriminate old stimuli from new stimuli. It is calculated as the difference between z-scored hit rates and false alarm rates. The criterion describes the systematic bias to classify a stimulus as old (“liberal” bias) or new (“conservative” bias). It is calculated as the average of z-scored hit and false alarm rates multiplied by -1.

### Statistical comparisons

We compared task-evoked pupil responses during encoding between stimuli that were later remembered and those that were forgotten during the free recall tasks or the recognition task using paired sample t-tests and by determining the corresponding Bayes factors (**Fig. 2**).

For assessing the effect of task-irrelevant sounds on pupil size and behavioral metrics, we applied repeated measures ANOVAs with the (no-) task-irrelevant sound condition (i.e. no sound vs. individual SOAs) either during encoding or during recognition as the within-subject (4-level) factor. We further scrutinized the relationships testing the differences in the variable of interest between the no-task-irrelevant sound condition and the respective SOA condition against zero by means of Bayesian t-tests (**Fig. 3-5**).

We tested the relationships between task-evoked pupil responses during the recognition task and recognition behavior (**Fig. S3**). For each participant, we computed the Spearman (rank order) correlation between each of the behavioral metrics and the bin-wise task-evoked pupil amplitudes. We also ran a second-order polynomial regression using the bin-wise average task-evoked pupil response amplitude as predictor of the behavioral metrics. The sample of correlation coefficients and regression weights was then compared against 0 with a paired-samples t-test.

## Author Contributions

Conceptualization: JH, JWdG, LS, THD

Experimental design: PVP, JH, THD

Data acquisition: PVP

Formal analysis: JH, PVP Writing—original draft: JH, THD

Writing—review and editing: PVP, JWdG, LS, JH, THD

Supervision: THD

Funding acquisition: LS, THD

## Data and code availability

The datasets generated and analyzed during the current study as well as analysis scripts will be made available upon publication.

## References

Ades, H. W., & Brookhart, J. M. (1950). The Central Auditory Pathway. Journal of neurophysiology, 13(3), 189–205. 10.1152/jn.1950.13.3.189

Bear, M. F., & Singer, W. (1986). Modulation of Visual Cortical Plasticity by Acetylcholine and Noradrenaline. Nature, 320(6058), 172–176. 10.1038/320172a0

Beerendonk, L., Mejías, J. F., Nuiten, S. A., de Gee, J. W., Fahrenfort, J. J., & van Gaal, S. (2024). A disinhibitory circuit mechanism explains a general principle of peak performance during midlevel arousal. Proceedings of the National Academy of Sciences of the United States of America, 121(5), e2312898121. 10.1073/pnas.2312898121

Beerendonk, L., Mejías, J. F., Nuiten, S. A., de Gee, J. W., Zantvoord, J. B., Fahrenfort, J. J., & van Gaal, S. (2025). Adaptive arousal regulation: Pharmacologically shifting the peak of the Yerkes-Dodson curve by catecholaminergic enhancement of arousal. Proceedings of the National Academy of Sciences of the United States of America, 122(28), e2419733122. 10.1073/pnas.2419733122

Bergt, A., Urai, A. E., Donner, T. H., & Schwabe, L. (2018). Reading memory formation from the eyes. The European journal of neuroscience, 47(12), 1525–1533. 10.1111/ejn.13984

Bliss, T. V. P., & Collingridge, G. L. (1993). A Synaptic Model of Memory - Long-Term Potentiation in the Hippocampus. Nature, 361(6407), 31–39. 10.1038/361031a0

Bouret, S., & Sara, S. J. (2005). Network reset: a simplified overarching theory of locus coeruleus noradrenaline function. Trends in Neurosciences, 28(11), 574–582. 10.1016/j.tins.2005.09.002

Brainard, D. H. (1997). The psychophysics toolbox. Spatial Vision, 10(4), 433–436. 10.1163/156856897×00357

Breton-Provencher, V., Drummond, G. T., Feng, J. S., Li, Y. L., & Sur, M. (2022). Spatiotemporal dynamics of noradrenaline during learned behaviour. Nature, 606(7915), 732–738. 10.1038/s41586-022-04782-2

Breton-Provencher, V., & Sur, M. (2019). Active control of arousal by a locus coeruleus GABAergic circuit. Nature Neuroscience, 22(2), 218–228. 10.1038/s41593-018-0305-z

Clark, A. (2015). Pillow (PIL Fork) Documentation. readthedocs. Retrieved November 8, 2025, from https://buildmedia.readthedocs.org/media/pdf/pillow/latest/pillow.pdf

Clewett, D., & Murty, V. P. (2019). Echoes of Emotions Past: How Neuromodulators Determine What We Recollect. eNeuro, 6(2). 10.1523/eneuro.0108-18.2019

Clewett, D., Sakaki, M., Huang, R., Nielsen, S. E., & Mather, M. (2017). Arousal amplifies biased competition between high and low priority memories more in women than in men: The role of elevated noradrenergic activity. Psychoneuroendocrinology, 80, 80–91. 10.1016/j.psyneuen.2017.02.022

Clewett, D. V., Huang, R., Velasco, R., Lee, T.-H., & Mather, M. (2018). Locus Coeruleus Activity Strengthens Prioritized Memories Under Arousal. The Journal of neuroscience : the official journal of the Society for Neuroscience, 38(6), 1558–1574. 10.1523/jneurosci.2097-17.2017

Cronin, S. L., Lipp, O. V., & Marinovic, W. (2023). Pupil dilation during encoding, but not type of auditory stimulation, predicts recognition success in face memory. Biological Psychology, 178, 108547. 10.1016/j.biopsycho.2023.108547

Dayan, P., & Yu, A. J. (2006). Phasic norepinephrine: A neural interrupt signal for unexpected events. Network-Computation in Neural Systems, 17(4), 335–350. 10.1080/09548980601004024

de Gee, J. W., Colizoli, O., Kloosterman, N. A., Knapen, T., Nieuwenhuis, S., & Donner, T. H. (2017). Dynamic modulation of decision biases by brainstem arousal systems. eLife, 6, e23232. 10.7554/eLife.23232

de Gee, J. W., Kloosterman, N. A., Braun, A., & Donner, T. H. (2025). Catecholamines reduce choice history biases in perceptual decision making. Plos Biology, 23(9), e3003361. 10.1371/journal.pbio.3003361

de Gee, J. W., Knapen, T., & Donner, T. H. (2014). Decision-related pupil dilation reflects upcoming choice and individual bias. Proceedings of the National Academy of Sciences of the United States of America, 111(5), E618–E625. 10.1073/pnas.1317557111

de Gee, J. W., Tsetsos, K., Schwabe, L., Urai, A. E., McCormick, D., McGinley, M. J., & Donner, T. H. (2020). Pupil-linked phasic arousal predicts a reduction of choice bias across species and decision domains. eLife, 9. 10.7554/eLife.54014

Dringenberg, H. C. (2020). The history of long-term potentiation as a memory mechanism: Controversies, confirmation, and some lessons to remember. Hippocampus, 30(9), 987–1012. 10.1002/hipo.23213

Goldinger, S. D., & Papesh, M. H. (2012). Pupil Dilation Reflects the Creation and Retrieval of Memories. Current Directions in Psychological Science, 21(2), 90–95. 10.1177/0963721412436811

Goto, A., Bota, A., Miya, K., Wang, J. B., Tsukamoto, S., Jiang, X. Z., Hirai, D., Murayama, M., Matsuda, T., McHugh, T. J., Nagai, T., & Hayashi, Y. (2021). Stepwise synaptic plasticity events drive the early phase of memory consolidation. Science, 374(6569), 857–863. 10.1126/science.abj9195

Grant, S. J., Aston-Jones, G., & Redmond, D. E. (1988). Responses of Primate Locus Coeruleus Neurons to Simple and Complex Sensory Stimuli. Brain Research Bulletin, 21(3), 401–410. 10.1016/0361-9230(88)90152-9

Green, D. M., & Swets, J. A. (1966). Signal detection theory and psychophysics (Vol. 1). New York: Wiley.

Hagena, H., & Manahan-Vaughan, D. (2025). Oppositional and competitive instigation of hippocampal synaptic plasticity by the VTA and locus coeruleus. Proceedings of the National Academy of Sciences of the United States of America, 122(1), e2402356122. 10.1073/pnas.2402356122

Harley, C. W. (2007). Norepinephrine and the dentate gyrus. Dentate Gyrus: A Comphrehensive Guide to Structure, Function, and Clinical Implications, 163, 299–318. 10.1016/S0079-6123(07)63018-0

Hebart, M. N., Dickter, A. H., Kidder, A., Kwok, W. Y., Corriveau, A., Van Wicklin, C., & Baker, C. (2019). THINGS: A database of 1,854 object concepts and more than 26,000 naturalistic object images. Plos One, 14(10), e0223792. 10.1371/journal.pone.0223792

Hebisch, J., Ghassemieh, A. C., Zhecheva, E., Brouwer, M., van Gaal, S., Schwabe, L., Donner, T. H., & de Gee, J. W. (2024). Task-irrelevant stimuli reliably boost phasic pupil-linked arousal but do not affect decision formation. Scientific Reports, 14(1), 28380. 10.1038/s41598-024-78791-8

Hoeks, B., & Levelt, W. J. M. (1993). Pupillary Dilation as a Measure of Attention - a Quantitative System-Analysis. Behavior Research Methods Instruments & Computers, 25(1), 16–26. 10.3758/Bf03204445

Jordan, R., & Keller, G. B. (2023). The locus coeruleus broadcasts prediction errors across the cortex to promote sensorimotor plasticity. eLife, 12. 10.7554/eLife.85111

Joshi, S., & Gold, J. I. (2020). Pupil Size as a Window on Neural Substrates of Cognition. Trends in Cognitive Sciences, 24(6), 466–480. 10.1016/j.tics.2020.03.005

Joshi, S., & Gold, J. I. (2022). Context-dependent relationships between locus coeruleus firing patterns and coordinated neural activity in the anterior cingulate cortex. eLife, 11. 10.7554/eLife.63490

Joshi, S., Li, Y., Kalwani, R. M., & Gold, J. I. (2016). Relationships between Pupil Diameter and Neuronal Activity in the Locus Coeruleus, Colliculi, and Cingulate Cortex. Neuron, 89(1), 221–234. 10.1016/j.neuron.2015.11.028

Knapen, T., de Gee, J. W., Brascamp, J., Nuiten, S., Hoppenbrouwers, S., & Theeuwes, J. (2016). Cognitive and Ocular Factors Jointly Determine Pupil Responses under Equiluminance. Plos One, 11(5), e0155574. 10.1371/journal.pone.0155574

Krishnamurthy, K., Nassar, M. R., Sarode, S., & Gold, J. I. (2017). Arousal-related adjustments of perceptual biases optimize perception in dynamic environments. Nature Human Behaviour, 1(6), 0107. 10.1038/s41562-017-0107

Kucewicz, M. T., Dolezal, J., Kremen, V., Berry, B. M., Miller, L. R., Magee, A. L., Fabian, V., & Worrell, G. A. (2018). Pupil size reflects successful encoding and recall of memory in humans. Scientific Reports, 8(1), 4949. 10.1038/s41598-018-23197-6

Lempert, K. M., Glimcher, P. W., & Phelps, E. A. (2015). Emotional Arousal and Discount Rate in Intertemporal Choice Are Reference Dependent. Journal of Experimental Psychology-General, 144(2), 366–373. 10.1037/xge0000047

Lewandowsky, S., Oberauer, K., & Brown, G. D. A. (2009). No temporal decay in verbal short-term memory. Trends in Cognitive Sciences, 13(3), 120–126. 10.1016/j.tics.2008.12.003

Macmillan, N. A., & Creelman, C. D. (2005). Detection Theory: A user’s Guide (2 ed.). Mahwah: Lawrence Erlbaum Associates.

Maheu, M., Donner, T. H., & Wiegert, J. S. (2025). bioRxiv, 2025–2003. 10.1101/2025.03.26.644382

Martin, S. J., Grimwood, P. D., & Morris, R. G. (2000). Synaptic plasticity and memory: an evaluation of the hypothesis. Annual review of neuroscience, 23(1), 649–711. 10.1146/annurev.neuro.23.1.649

Mather, M., & Sutherland, M. R. (2011). Arousal-Biased Competition in Perception and Memory. Perspectives on Psychological Science, 6(2), 114–133. 10.1177/1745691611400234

McGinley, M. J., David, S. V., & McCormick, D. A. (2015). Cortical Membrane Potential Signature of Optimal States for Sensory Signal Detection. Neuron, 87(1), 179–192. 10.1016/j.neuron.2015.05.038

Nassar, M. R., Rumsey, K. M., Wilson, R. C., Parikh, K., Heasly, B., & Gold, J. I. (2012). Rational regulation of learning dynamics by pupil-linked arousal systems. Nature Neuroscience, 15(7), 1040–1046. 10.1038/nn.3130

Nuiten, S., de Gee, J. W., Zantvoord, J., Sterzer, P., Fahrenfort, J., & Gaal, S. v. (2025). Phasic and tonic arousal distinctly shape human decision bias. PREPRINT (Version 1) available at Research Square. 10.21203/rs.3.rs-6479550/v1

Petersen, A., Petersen, A. H., Bundesen, C., Vangkilde, S., & Habekost, T. (2017). The effect of phasic auditory alerting on visual perception. Cognition, 165, 73–81. 10.1016/j.cognition.2017.04.004

Poe, G. R., Foote, S., Eschenko, O., Johansen, J. P., Bouret, S., Aston-Jones, G., Harley, C. W., Manahan-Vaughan, D., Weinshenker, D., Valentino, R., Berridge, C., Chandler, D. J., Waterhouse, B., & Sara, S. J. (2020). Locus coeruleus: a new look at the blue spot. Nature Reviews Neuroscience, 21(11), 644–659. 10.1038/s41583-020-0360-9

Ratcliff, R. (1978). A Theory of Memory Retrieval. Psychological review, 85(2), 59–108.

Richter-Levin, G., & Akirav, I. (2003). Emotional tagging of memory formation – in the search for neural mechanisms. Brain research. Brain research reviews, 43(3), 247–256. 10.1016/j.brainresrev.2003.08.005

Schacter, D. L., & Addis, D. R. (2009). Remembering the Past to Imagine the Future: A Cognitive Neuroscience Perspective. Military Psychology, 21(sup1), S108–S112. 10.1080/08995600802554748

Schwarz, L. A., & Luo, L. Q. (2015). Organization of the Locus Coeruleus-Norepinephrine System. Current Biology, 25(21), R1051–R1056. 10.1016/j.cub.2015.09.039

Seol, G. H., Ziburkus, J., Huang, S., Song, L., Kim, I. T., Takamiya, K., Huganir, R. L., Lee, H. K., & Kirkwood, A. (2007). Neuromodulators control the polarity of spike-timing-dependent synaptic plasticity. Neuron, 55(6), 919–929. 10.1016/j.neuron.2007.08.013

Shadlen, M. N., & Shohamy, D. (2016). Decision Making and Sequential Sampling from Memory. Neuron, 90(5), 927–939. 10.1016/j.neuron.2016.04.036

Shohamy, D., & Adcock, R. A. (2010). Dopamine and adaptive memory. Trends in Cognitive Sciences, 14(10), 464–472. 10.1016/j.tics.2010.08.002

Sonneborn, A., & Greene, R. W. (2021). Norepinephrine transporter antagonism prevents dopamine-dependent synaptic plasticity in the mouse dorsal hippocampus. Neuroscience Letters, 740(135450). 10.1016/j.neulet.2020.135450

Tambini, A., Rimmele, U., Phelps, E. A., & Davachi, L. (2017). Emotional brain states carry over and enhance future memory formation. Nature Neuroscience, 20(2), 271–278. 10.1038/nn.4468

Teigen, K. H. (1994). Yerkes-Dodson - a Law for All Seasons. Theory & Psychology, 4(4), 525–547. 10.1177/0959354394044004

Tona, K. D., Murphy, P. R., Brown, S. B. R. E., & Nieuwenhuis, S. (2016). The accessory stimulus effect is mediated by phasic arousal: A pupillometry study. Psychophysiology, 53(7), 1108–1113. 10.1111/psyp.12653

Totah, N. K., Neves, R. M., Panzeri, S., Logothetis, N. K., & Eschenko, O. (2018). The Locus Coeruleus Is a Complex and Differentiated Neuromodulatory System. Neuron, 99(5), 1055–1068. 10.1016/j.neuron.2018.07.037

van den Brink, R. L., Murphy, P. R., & Nieuwenhuis, S. (2016). Pupil Diameter Tracks Lapses of Attention. Plos One, 11(10), e0165274. 10.1371/journal.pone.0165274

Wang, C. A., & Munoz, D. P. (2014). Modulation of stimulus contrast on the human pupil orienting response. European Journal of Neuroscience, 40(5), 2822–2832. 10.1111/ejn.12641

Yerkes, R. M., & Dodson, J. D. (1908). The relation of strength of stimulus to rapidity of habit-formation. Journal of Comparative Neurology and Psychology, 18, 459–482.

