## Supplemental Figures for "Task-irrelevant stimuli boost phasic pupil-linked arousal but not memory formation"

[1]Section Computational Cognitive Neuroscience, Department of Neurophysiology and Pathophysiology, University Medical Center Hamburg-Eppendorf, Hamburg, DEU; [2]Zurich Center for Neuroeconomics (ZNE), Department of Economics, University of Zurich, Zurich, CH; [3]University Research Priority Program "Adaptive Brain Network Mechanisms in Development and Learning" (URPP AdaBD), University of Zurich, Zurich, CH; [4]Department of Cognitive Psychology, Institute of Psychology, University of Hamburg, Hamburg, DEU; [5]Cognitive and Systems Neuroscience, Swammerdam Institute for Life Sciences, University of Amsterdam, Amsterdam, NLD; [6]Amsterdam Brain & Cognition, University of Amsterdam, Amsterdam, NLD; [7]Bernstein Center for Computational Neuroscience, Charité Universitätsmedizin, Berlin, DEU.

### = Equal contribution

#### SUPPLEMENTARY FIGURES

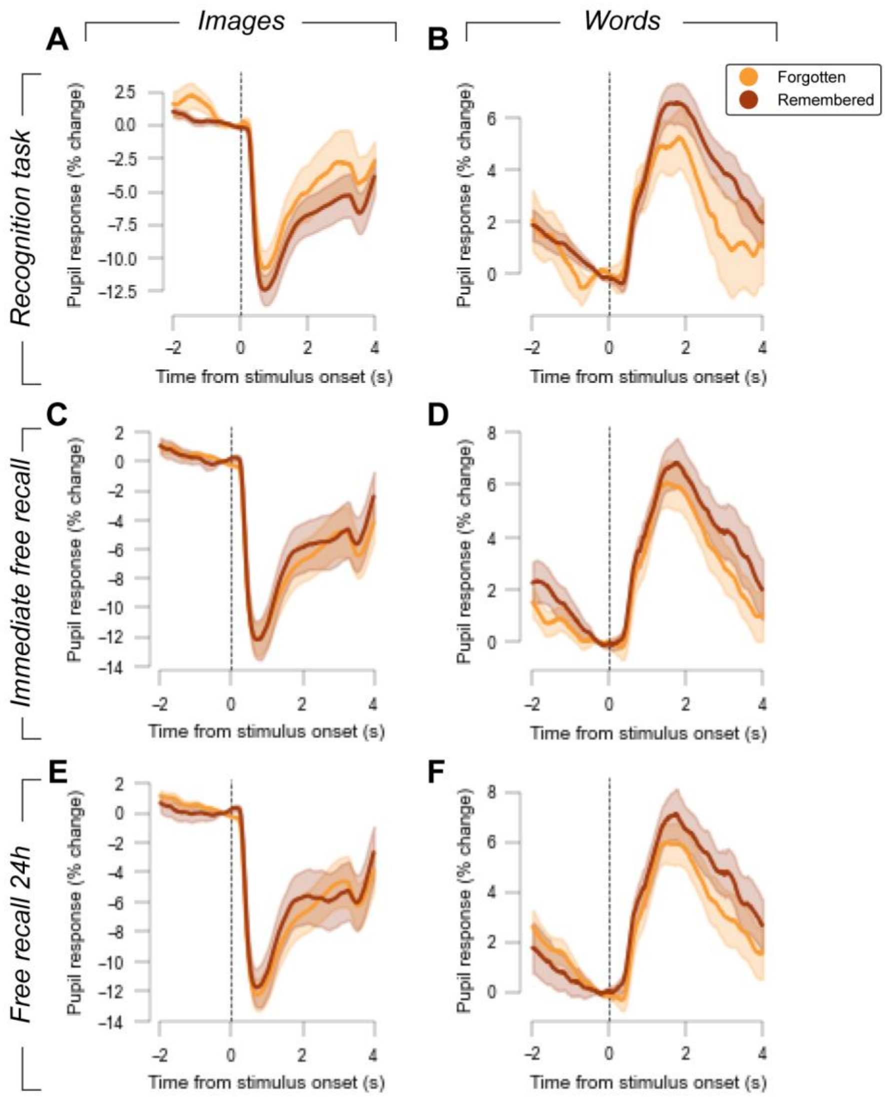


**Figure S1. Pupil responses during encoding sorted by subsequent memory performance. (A)** Pupil response time course during encoding on trials with no task-irrelevant sound for images that were subsequently recognized (remembered) vs. missed (forgotten) in the recognition task. Shading, SEM across participants. **(B)** As (A), but for the word encoding task. **(C-D)** As (A-B), but for images and words that were immediately recalled vs. not recalled after the encoding task. **(E-F)** As (A-B) and (C-D), but for images and words that were recalled vs. not recalled 24h later.

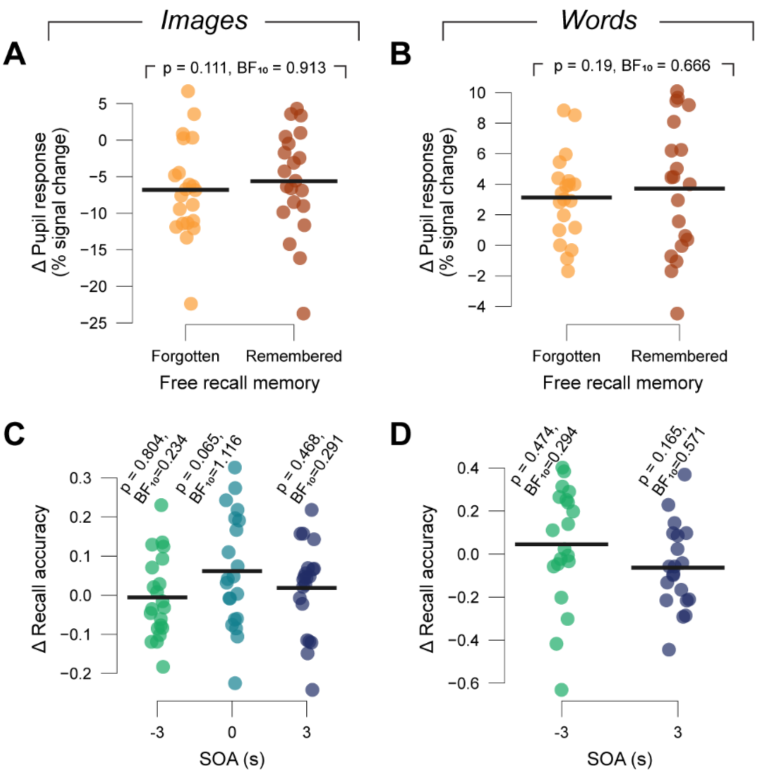


**Figure S2. Effects of pupil response and task-irrelevant sounds on immediate free recall.** **(A)** Individual trial-average pupil response amplitudes during encoding (collapsed across an interval of 1-3 s from image onset) on trials with no task-irrelevant sounds. Trials are sorted by whether images were recalled (remembered) or not (forgotten) in the free recall task on day 1. Black bars, group average. No increase in pupil response was observed for remembered versus forgotten images: t=1.261, p=0.111, BF10=0.913, one-sided t-test. **(B)** Same as (A) but for the word task. Again, no evidence was found for a group effect: t=0.901, p=0.19, BF10=0.666, one-sided t-test. **(C)** Difference in accuracy of day 1 recall between trials with and without task-irrelevant sounds for each participant, sorted by sound SOA during encoding. We found no main effect of the sound: $F_{(3,60)}$=1.918, p=0.136. Black bars, group average. P and $BF_{10}$, p-values and Bayes factors for t-tests against zero. **(D)** Same as (C) but for the word task. Again, there was no effect of task-irrelevant sounds: $F_{(2,38)}$=1.655, p=0.205.

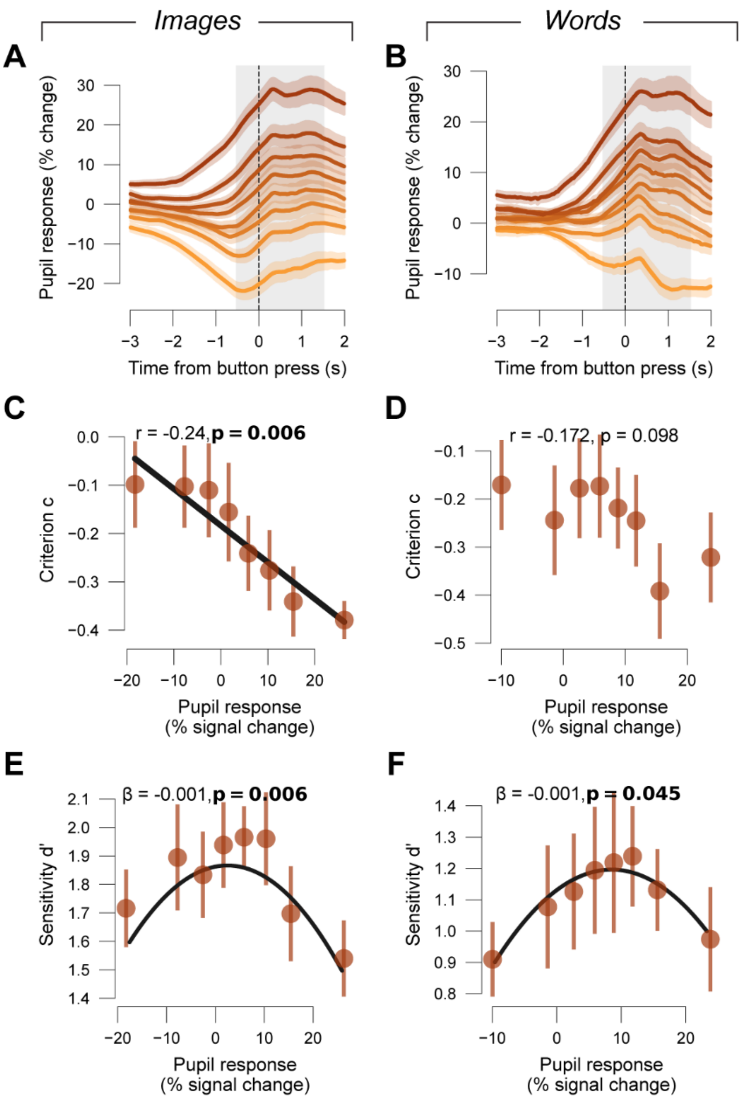


**Figure S3. Link between pupil response amplitudes and behavior during recognition task. (A-B)** Pupil response time course in the Recognition task on trials without task-irrelevant sounds, locked to decision button press, binned post-hoc by size of task-evoked pupil response. Shading, SEM across participants. Grey shading, interval used for quantifying task-evoked pupil responses (see Methods). **(C-D)** Decision bias (criterion c) in dependance of task-evoked pupil response in the Recognition task on trials without task-irrelevant sounds. Error bars, S.E.M. across participants. R-values, Spearman correlation. **(E-F)** As (C-D), but for decision performance (sensitivity d'). β, weight of 2$^{nd}$ degree polynomial regression. Statistics for image task: group average coefficient of the quadratic effect: β=-0.001, t=-3.038, p=0.006; Statistics for word task group average coefficient of the quadratic effect: β=-0.001, t=-2.15, p<0.045 (polynomial regression).
